# Pacybara: Accurate long-read sequencing for barcoded mutagenized allelic libraries

**DOI:** 10.1101/2023.02.22.529427

**Authors:** Jochen Weile, Gabrielle Ferra, Gabriel Boyle, Sriram Pendyala, Clara Amorosi, Chiann-Ling Yeh, Atina G. Cote, Nishka Kishore, Daniel Tabet, Warren van Loggerenberg, Ashyad Rayhan, Douglas M Fowler, Maitreya J. Dunham, Frederick P Roth

## Abstract

**Summary:** Long read sequencing technologies, an attractive solution for many applications, often suffer from higher error rates. Alignment of multiple reads can improve base-calling accuracy, but some applications, e.g. sequencing mutagenized libraries where multiple distinct clones differ by one or few variants, require the use of barcodes or unique molecular identifiers. Unfortunately, sequencing errors can interfere with correct barcode identification, and a given barcode sequence may be linked to multiple independent clones within a given library.

Here we focus on the target application of sequencing mutagenized libraries in the context of multiplexed assays of variant effects (MAVEs). MAVEs are increasingly used to create comprehensive genotype-phenotype maps that can aid clinical variant interpretation. Many MAVE methods use long-read sequencing of barcoded mutant libraries for accurate association of barcode with genotype. Existing long-read sequencing pipelines do not account for inaccurate sequencing or non-unique barcodes. Here, we describe Pacybara, which handles these issues by clustering long reads based on the similarities of (error-prone) barcodes while also detecting barcodes that have been associated with multiple genotypes. Pacybara also detects recombinant (chimeric) clones and reduces false positive indel calls. In three example applications, we show that Pacybara identifies and correctly resolves these issues.

**Availability and Implementation:** Pacybara, freely available at https://github.com/rothlab/pacybara, is implemented using R, Python and bash for Linux. It has both a single-threaded implementation and, for GNU/Linux clusters that use Slurm, PBS, or GridEngine schedulers, a multi-node version.

**Supplementary Material:** Supplementary materials are available at Bioinformatics online.

## Introduction

Multiplexed Assays of Variant Effect (MAVEs) often involve the use of clone libraries in which each mutagenized allele is associated with a barcode (Tabet et al. 2022), requiring determination of full-length amplicon sequences. Long-read sequencing technologies such as Pacbio Sequel II, Revio and Oxford Nanopore have become an attractive alternative to short-read assembly methods (Hiatt et al. 2010; Weile et al. 2017).

The first pipeline for long-read barcoded library assembly was AssemblyByPacbio (ABP) (Matreyek et al. 2018). ABP extracts barcode sequences from each read, groups reads by identical barcodes, and derives the clone genotype from the highest quality read in each group. Recently, the PacRAT pipeline was developed in which the ABP output is post-processed to derive a multiple sequence alignment-based consensus for each read group sharing a barcode (Yeh & Amorosi et al. 2022).

At least three problems have remained unsolved: First, sequencing errors in the barcode can cause a single clone to appear as two or more distinct clones. Second, a barcode sequence may be “non-unique”, in that the same sequence is associated with multiple clones of different genotypes within a given library. Although an error-aware clustering method for short barcode sequences (Zhao et al. 2018) addresses the first issue for short read barcode identification, it does not consider clonality of genotypes beyond the barcode itself. Third, problems arising during library preparation, e.g. PCR crossover events, can yield consensus genotypes that are recombinant chimeras of wild-type (WT) or other variant genotypes. Karst and colleagues have described a double-barcode method to detect chimeras, but this approach is not applicable to singly-barcoded libraries in which clones might differ by only a few SNVs, as is the case for mutagenized libraries used in MAVEs (Karst et al. 2021).

To address these issues, we extended the clustering approaches cited above using not only the barcode sequences from each read but also the full set of candidate variants and their quality metrics. Although initial stages of the clustering process, like previous methods, focused on merging pairs of reads with identical candidate barcodes, Pacybara accounts for variant quality and the extent of overlap between candidate variants. Pacybara then further considers merging of reads with similar but non-identical barcodes.

## Methods

Pacybara, designed to run on high-performance computing clusters, consists of a main executable that deploys and supervises individual jobs on cluster nodes and collates the results. In the first processing step, Pacbio HiFi reads are distributed across jobs and aligned to the reference sequence. From the alignments, barcode sequences and lists of candidate variants (i.e. basecalls that apparently differ from the reference sequence) are extracted, including their respective quality scores. For libraries designed with multiple barcodes per molecule, the latter can optionally be combined to form a single “virtual barcode” each. Reads in which quality scores within the barcode region fall below a tunable threshold (default Q62) are filtered out, as they often indicate multi-occupancy SMRT wells.

Clustering first considers sets of reads with identical barcode sequences. Within each identical-barcode set, putative sequencing errors are excluded by filtering out candidate variants seen in only one read and with a below-threshold quality score. Then for each identical-barcode read set, a graph is constructed such that each node corresponds to a read, and an edge is added between two reads either if both are WT or if they harbor a sufficiently similar set of candidate variants (defined by a Jaccard index threshold). A “seed cluster” is then formed for each graph component (including ‘singleton’ nodes without edges).

Cluster definitions, initially defined by the set of seed clusters, are updated in a cluster-merging process. Pairs of clusters, chosen by the minimum edit distance (ED) of their respective member reads, are iteratively considered as candidate clusters up to a maximum edit distance (default 2). After filtering sequencing errors from this candidate cluster as above, the candidate cluster is accepted if all of the following are true: a) all reads in both clusters are WT or the sets of remaining candidate variants in the respective pair of clusters have an above-threshold Jaccard coefficient, b) no pair of reads in the proposed cluster has a barcode edit distance greater than the maximally allowed distance, and c) the pair of clusters has sufficiently divergent sizes, as might be expected if one cluster represents a subset of reads derived from the same clone, but with more base-calling errors in the barcode. Here the clusters were considered to have sufficiently divergent sizes if |log2(size_1_/size_2_)| > ED, similar to a previous approach for short-read barcode clustering (Zhao et al. 2018). We used a Jaccard threshold of 0.2 in each case.

For each cluster in the final set, consensus barcodes are derived, variants are filtered as above to remove putative base-calling errors, and (for coding sequences) remaining variants are translated to protein consequences.

## Results and Discussion

We evaluated performance of both Pacybara and the previously-established method PacRAT. Both tools were applied to three different barcoded mutagenized open reading frame libraries, designed for use in a MAVE: (1) A mutagenized human *CYP2C9* library (Amorosi et al. 2021), (2) a mutagenized human *CYP2C19* library (Boyle et al. 2023), and (3) a previously unpublished mutagenized human *LDLR* library, which we suspected of harboring non-unique barcodes and PCR chimeras.

Differences between Pacybara and PacRAT were immediately apparent in the distribution of cluster sizes (Figure S1). In all three datasets, Pacybara produced a larger number of small clusters, which we attribute to (i) more aggressive filtering of low-quality reads and (ii) fewer erroneous merges between clones with non-unique barcodes by considering the coherence of clone genotypes before merging.

For all three datasets, both pipelines reported similar numbers of clusters and similar distributions of the number of mutations. A notable exception to this is frameshift variants in *CYP2C9*, nearly half of which were filtered out by Pacybara (Figure S2). This is consistent with false-positive indels being a common type of sequencing error in PacBio sequencing, especially at homopolymer sites. Indeed, we found that frameshift calls at the 5’ end of homopolymer runs of length 4 or greater were 2.3 times as likely to be filtered out by Pacybara compared to other positions (Figure S3).

We next wished to examine how well variant calls agreed between PacRAT and Pacybara for each barcoded clone. Of those clones that were not filtered out by either tool, 88%, 79%, and 96% agreed completely (Figure S4). We then examined the raw read data of five arbitrarily chosen clusters with disagreeing genotypes. In three out of five cases, the pipelines primarily disagreed on the exact lengths of long indels. The remaining two cases disagreed on whether to include or exclude low-quality variants that only occur in some but not all reads within a cluster (see Supplementary Text S1).

To validate whether the barcode-genotype associations produced by either pipeline are more accurate, we tested them in terms of their impact on protein function. As nonsense and frameshift variants are likely to damage protein function while synonymous variants are not, we compared, for each pipeline, the distributions of functionality scores for clones assigned a nonsense and frameshift variant against those bearing only synonymous variants. Functional scores were calculated based on enrichment or depletion in a fluorescence-based assay (measuring protein abundance for CYP2C19, enzyme activity for CYP2C9, and the ability of the cell to uptake LDL particles for LDLR). For all three data sets, score distributions based on both pipelines were very similar, with Pacybara’s frameshift scores being slightly more likely to be deleterious. (Figure S5).

Next, we examined whether Pacybara was able to detect clones with non-unique barcodes. For the *CYP2C9* and *CYP2C19* libraries, only a small fraction of barcodes were non-unique (Figure S6). For the *LDLR* library 22.4% of barcodes were non-unique. However, 17.3% were classified as “remediable”, in that a single clone represented more than two thirds of reads for that barcode and would thus likely dominate readouts for that barcode in a screen. We manually examined eight arbitrarily chosen non-unique barcodes from the *LDLR* library which had conflicting genotypes assigned by the two pipelines. In one case, PacRAT called the genotype correctly, while Pacybara did not, as it included a frequently occurring sequencing error occurring in multiple reads. However, in six out of the eight cases Pacybara called the correct genotype by splitting the cluster. One final case could not be conclusively decided due to ambiguous read data. (See Supplementary Text S2)

Pacybara’s use of virtual barcodes, concatenating barcodes from both flanks of the same construct, allows it to identify putative PCR crossover events. The *LDLR* library made use of such flanking barcodes, allowing us to test this feature. Indeed, after enabling virtual barcodes we identified 74,485 sets of clusters in which upstream barcodes were identical and either complete or partial overlap in genotype existed, but which had entirely different downstream barcodes (Table S1). We consider these cases likely PCR chimeras. To investigate the relationship between the putative chimeras and barcodes that are found to be non-unique without the use of virtual barcodes, we counted how often single upstream barcodes were flagged as a chimera or non-unique or both. We found that for 99.99% of cases in which a barcode was flagged as non-unique, it was also found to be involved in a putative PCR chimera. Similarly, if a barcode was not involved in a chimera, it had a 99.99% chance of being unique. However, out of all barcodes putatively involved in chimeras, it had only a 21% chance of being detected as non-unique when virtual barcodes were disabled. This observation confirms the value of dual flanking barcodes, which were unfortunately not a feature of the *CYP2C9* and *CYP2C19* libraries, while showing that detecting non-unique barcodes can at least serve as a way of alleviating problems caused by chimeras. No other pipeline has been described that detects chimeras in barcoded clone libraries where clones differ by only one or few nucleotides.

In summary, we developed Pacybara to process long-read data from libraries of barcoded mutagenized clones, and found that it successfully detects non-unique barcodes and PCR crossover events, combines reads with barcodes that differ only due to sequencing error, reduces false-positive indel calls, and thus improves the quality of a downstream MAVE experiment.

## Acknowledgements

We would like to acknowledge Kelley Harris for her advice. We also acknowledge funding from the National Human Genome Research Institute (NHGRI) of the National Institutes of Health (NIH) Center of Excellence in Genomic Science Initiative (CEGS) (RM1 HG010461), the Center for Actionable Variant Analysis (UM1 HG011969), the NIH R01 HL52066, and the NHGRI Impact of Genomic Variation on Function Initiative (UM1 HG011989) to D.M.F, a National Institute of General Medical Sciences of the NIH grant (R01 GM132162) to M.J.D and D.M.F., the NIH R01 HL164675 to F.P.R, and a Canadian Institutes of Health Research Foundation Grant to F.P.R.

## References

Amorosi, Clara J., Melissa A. Chiasson, Matthew G. McDonald, Lai Hong Wong, Katherine A. Sitko, Gabriel Boyle, John P. Kowalski, Allan E. Rettie, Douglas M. Fowler, and Maitreya J. Dunham. 2021. “Massively Parallel Characterization of CYP2C9 Variant Enzyme Activity and Abundance.” American Journal of Human Genetics 108 (9): 1735–51.

Boyle, Gabriel E., Katherine Sitko, Jared G. Galloway, Hugh K. Haddox, Aisha Haley Bianchi, Ajeya Dixon, Raine E. S. Thomson, et al. 2023. “Deep Mutational Scanning of CYP2C19 Reveals a Substrate Specificity-Abundance Tradeoff.” bioRxiv. 10.1101/2023.10.06.561250.

Hiatt, Joseph B., Rupali P. Patwardhan, Emily H. Turner, Choli Lee, and Jay Shendure. 2010. “Parallel, Tag-Directed Assembly of Locally Derived Short Sequence Reads.” Nature Methods. 10.1038/nmeth.1416.

Karst, Søren M., Ryan M. Ziels, Rasmus H. Kirkegaard, Emil A. Sørensen, Daniel McDonald, Qiyun Zhu, Rob Knight, and Mads Albertsen. 2021. “High-Accuracy Long-Read Amplicon Sequences Using Unique Molecular Identifiers with Nanopore or PacBio Sequencing.” Nature Methods 18 (2): 165–69.

Matreyek, Kenneth A., Lea M. Starita, Jason J. Stephany, Beth Martin, Melissa A. Chiasson, Vanessa E. Gray, Martin Kircher, et al. 2018. “Multiplex Assessment of Protein Variant Abundance by Massively Parallel Sequencing.” Nature Genetics 50 (6): 874–82.

Tabet, Daniel, Victoria Parikh, Prashant Mali, Frederick P. Roth, and Melina Claussnitzer. 2022. “Scalable Functional Assays for the Interpretation of Human Genetic Variation.” Annual Review of Genetics, September. 10.1146/annurev-genet-072920-032107.

Weile, Jochen, Song Sun, Atina G. Cote, Jennifer Knapp, Marta Verby, Joseph C. Mellor, Yingzhou Wu, et al. 2017. “A Framework for Exhaustively Mapping Functional Missense Variants.” Molecular Systems Biology 13 (12): 957.

Yeh, Chiann-Ling C., Clara J. Amorosi, Soyeon Showman, and Maitreya J. Dunham. 2022. “PacRAT: A Program to Improve Barcode-Variant Mapping from PacBio Long Reads Using Multiple Sequence Alignment.” Bioinformatics 38 (10): 2927–29.

Zhao, Lu, Zhimin Liu, Sasha F. Levy, and Song Wu. 2018. “Bartender: A Fast and Accurate Clustering Algorithm to Count Barcode Reads.” Bioinformatics 34 (5): 739–47.

