## Supplementary Materials for "Pacybara: Accurate long-read sequencing for barcoded mutagenized allelic libraries"

### Supplementary Figures

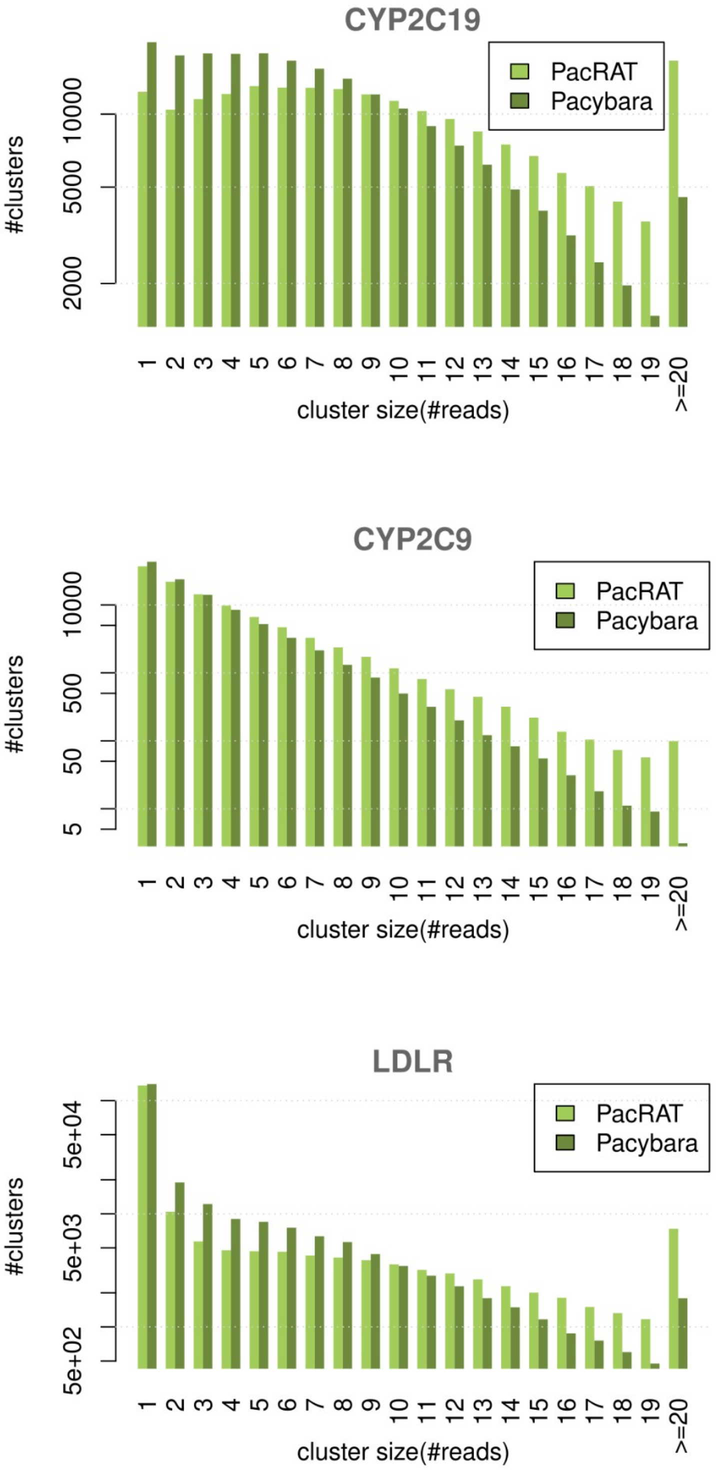

Figure S1: A: Comparison of cluster size distributions. Both PacRAT and Pacybara were applied to the same datasets (mutagenized library of *CYP2C9*, *CYP2C19*, and *LDLR*).

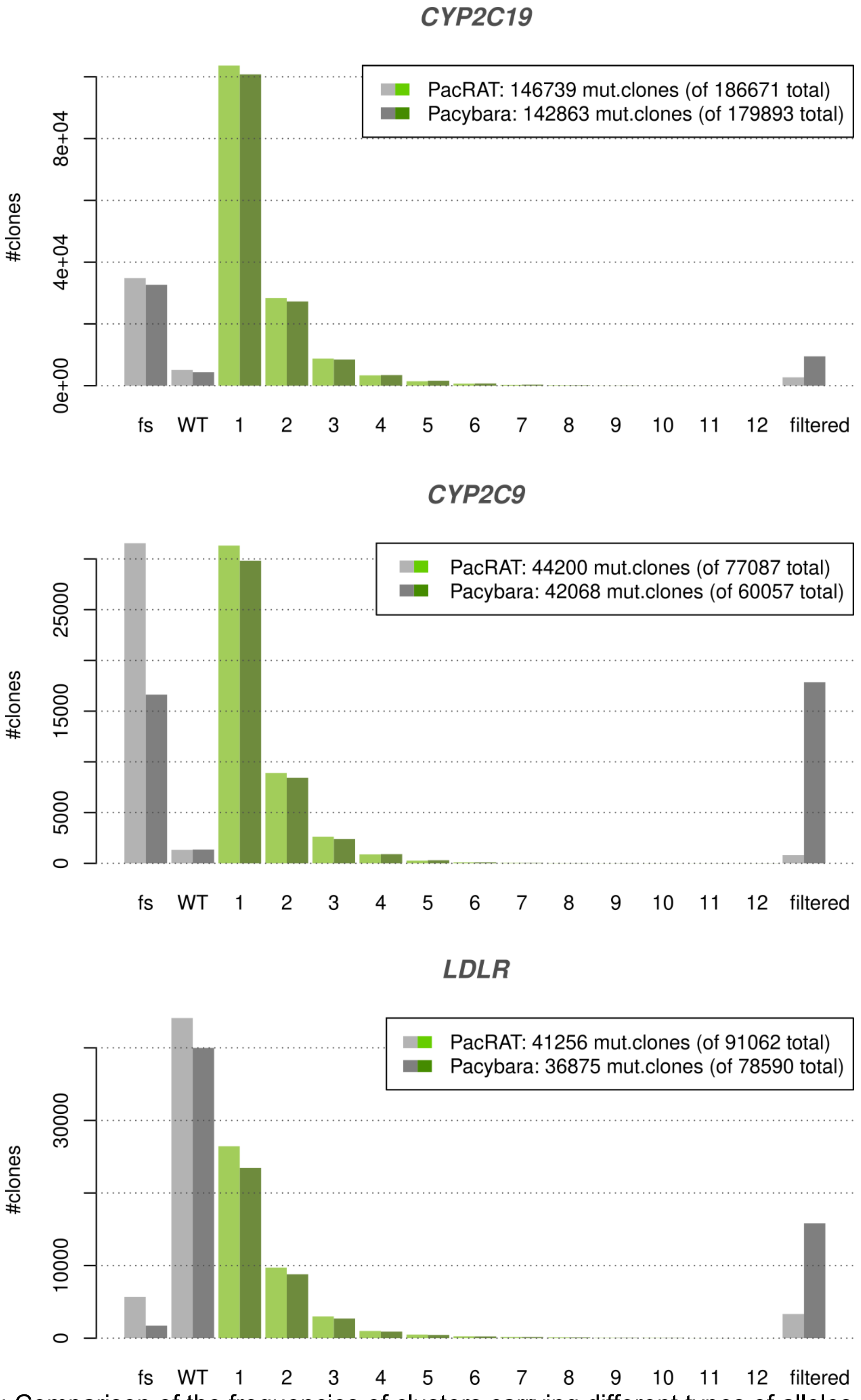

Figure S2: Comparison of the frequencies of clusters carrying different types of alleles. fs = frameshift; WT = wildtype allele; 1-12 = carrying the corresponding number of codon changes; filtered = clusters filtered out for low quality.

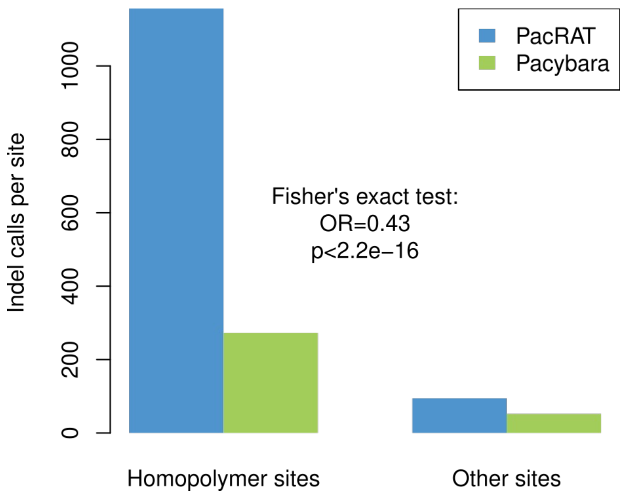

Figure S3: Indel/Frameshift calls made by PacRAT and Pacybara at homopolymer sites and all other sites within the *CYP2C9* sequence.

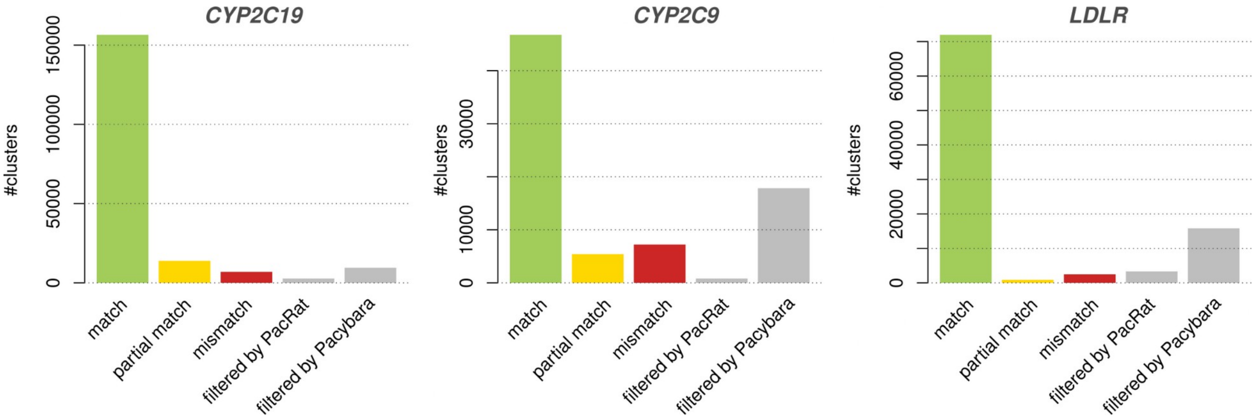

Figure S4: Comparison of cluster variant calls between PacRAT and Pacybara. Match: Both tools agreed on genotype, Partial match: Both tools agreed at least one, but not all nucleotide changes; Mismatch: Both tools disagree on the genotype; Filtered: Clone was filtered out by given tool.

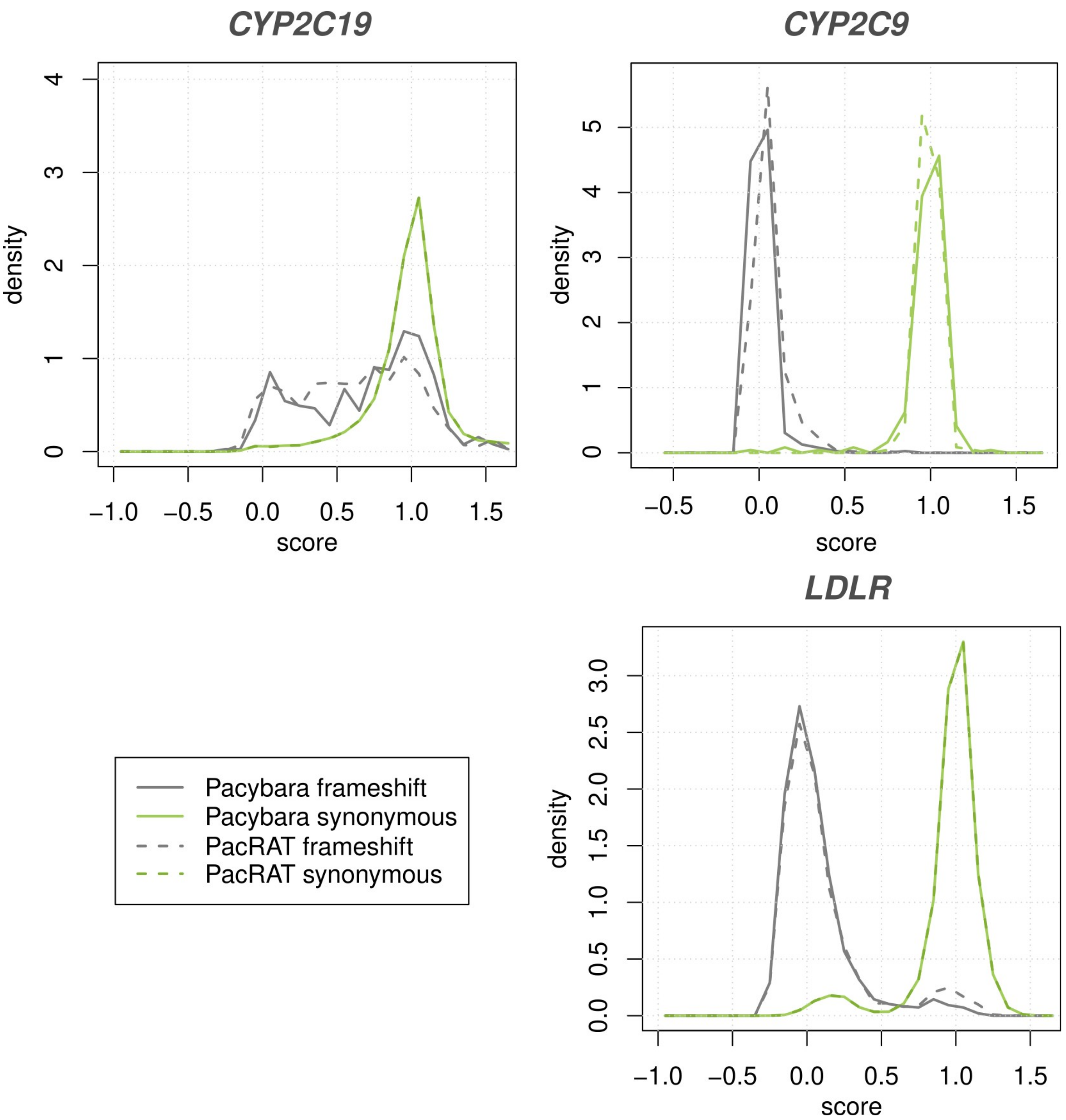

Figure S5: Distributions of functionality scores assigned to clones (i.e. clusters) identified by PacRAT and Pacybara respectively. Functionality scores act as a proxy for protein function. Specifically, for *CYP2C9* and *CYP2C19*, functionality scores represent regression coefficients across enrichment/depletion ratios of variants across fluorescence levels at different timepoints. For *LDLR*, functionality scores represent log-ratio of variant frequencies at high fluorescence levels over overall frequencies in the complete cell pool. Scores were normalized such that values near 1 indicate normal wildtype protein function, while scores near zero scores indicate complete loss of protein function.

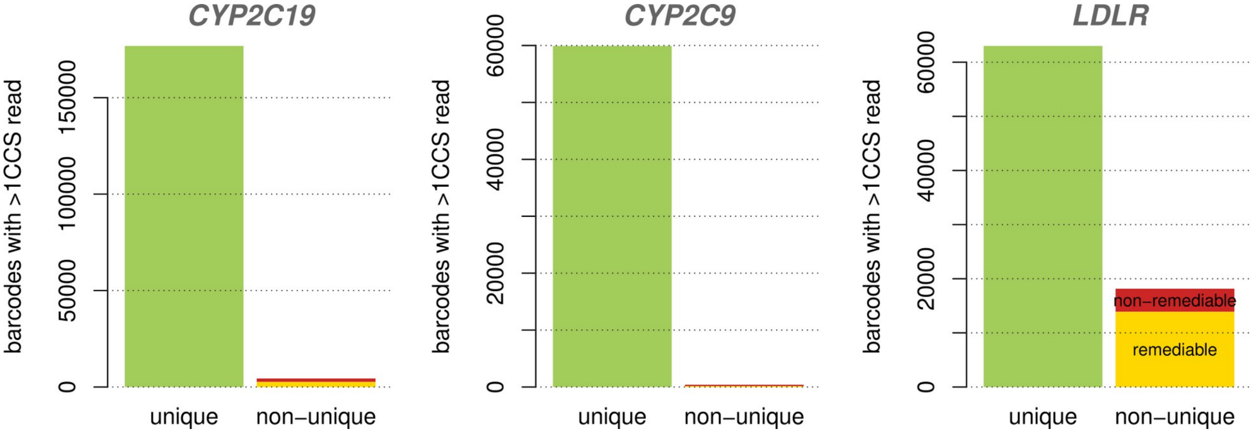

Figure S6: Breakdown of unique and non-unique barcodes assigned to clusters by Pacybara. The "remediable" category refers to clones containing greater than two thirds of reads for their barcode, which can be expected to dominate measurements of the barcode in a screen.

### Supplemental Text 1: Disagreeing variant call

Here we examined five arbitrarily chosen barcodes for which the pipelines disagreed on variant calls. In four out of five cases, the underlying alignment tools disagreed on length and/or positioning of long indels. In the remaining case, PacRAT included "frequent-flier" indel errors that only occurred in less than half of the underlying reads.

#### Barcode CGGCATGTGTTCGGGCAT

Pacybara: `c.[798T>C;799_801del] = p.[Cys266=;Phe267del]`

- 103del:93;798del:93;799del:93;800del:93;1258del:93;1361del:93
- 798del:93;799del:93;800del:93
- 798del:93;799del:93;800del:93
- 798del:93;799del:93;800del:93;1258del:57
- 798del:93;799del:93;800del:93;1258del:48
- 472del:58;798del:93;799del:93;800del:93
- 737del:54;798del:77;799del:77;800del:77;1258del:48
- 798del:93;799del:93;800del:93
- 381del:52;608del:51;798del:93;799del:93;800del:93;1598G>A:51

PacRAT: `c.796_798del;1258del = p.Lys420fs`

- 796del;797del;798del
- 796del;797del;798del
- 796del;797del;798del;1258del
- 796del;797del;798del;1258del
- 472del;796del;797del;798del
- 737del;796del;797del;798del;1258del
- 381del;608del;796del;797del;798del
- 103del;796del;797del;798del;1258del;1361del
- 796del;797del;798del

Here, PacRAT decided to include a variant that only occurred in 4 out of 9 reads, which resulted in a frameshift call.

#### Barcode TGGGTTCGAGGTGAGTTG

Pacybara: `c.[513_514insA;525_534del] = p.Gly171fs`

- 514insA:93;525del:93;526del:93;527del:93;528del:93;529del:93;530del:93;531del:93;532del:93;533del:93;534del:93
- -2del:49;193del:58;514insA:93;525del:86;526del:86;527del:86;528del:86;529del:86;530del:86;531del:86;532del:86;533del:86;534del:86;545del:48;678del:32;737del:48

- 88del:93;514insA:93;525del:93;526del:93;527del:93;528del:93;529del:93;530del:93;531del:93;532del:93;533del:93;534del:93;588del:93;1469del:82;1592del:93
- 514insA:93;525del:93;526del:93;527del:93;528del:93;529del:93;530del:93;531del:93;532del:93;533del:93;534del:93
- 514insA:93;525del:93;526del:93;527del:93;528del:93;529del:93;530del:93;531del:93;532del:93;533del:93;534del:93
- 514insA:93;525del:93;526del:93;527del:93;528del:93;529del:93;530del:93;531del:93;532del:93;533del:93;534del:93

PacRAT: `"c.514_525delinsATG = p.Cys172_Cys175delinsMet`

- 193del;513_514insA;516del;517del;518del;519del;520del;521del;522del;523del;524del;525del;545del;678del;737del
- 513_514insA;516del;517del;518del;519del;520del;521del;522del;523del;524del;525del
- 513_514insA;516del;517del;518del;519del;520del;521del;522del;523del;524del;525del
- 338del;513_514insA;516del;517del;518del;519del;520del;521del;522del;523del;524del;525del;690_691insA;1249del;1293del
- 88del;513_514insA;516del;517del;518del;519del;520del;521del;522del;523del;524del;525del;588del;1470del
- 30del;82del;87_88insC;87_88insA;92C>G;107_108insG;127del;142del;224_225insG;248del;267del;296del;318del;338del;345_346insA;429T>G;432del;501T>G;513_514insA;516del;517del;518del;519del;520del;521del;522del;523del;524del;525del;545del;588del;596A>C;621_622insC;652del;665del;747_748insG;840_841insG;846T>A;847A>C;903_904insA;959_960insG;966del;993_994insT;1014del;1015del;1016T>A;1026_1027insT;1191_1192insA;1219del;1227_1228insC;1230T>G;1293del;1358del;1362A>C;1394_1395insT
- 513_514insA;516del;517del;518del;519del;520del;521del;522del;523del;524del;525del
- 513_514insA;516del;517del;518del;519del;520del;521del;522del;523del;524del;525del

This is a disagreement on the length/position of the indel

#### Barcode TCTGTATCGCTATCATTG

Pacybara: `c.1462_1464delinsTGT = p.Ile488Cys`

- 1258del:48;1462A>T:88;1463T>G:84;1464A>T:93
- 1361del:59;1462A>T:93;1463T>G:93;1464A>T:93;1592del:52;1598G>A:16

PacRAT: `c.[1361_1361del;1462_1464delinsTGT] = "p.Gln454fs`

- 1258del;1462A>T;1463T>G;1464A>T
- 1361del;1462A>T;1463T>G;1464A>T

PacRAT got it wrong by including one of the random deletions

#### Barcode TATCCTAACCGAATATGA

Pacybara: `c.= = p.=`

- 1258del:48;1473del:81;1474del:81;1475del:81;1476del:81;1477del:81;1478del:81;1479del:81;1480del:81;1481del:81;1599A>G:61
- 1473del:93;1474del:93;1475del:93;1476del:93;1477del:93;1478del:93;1479del:93;1480del:93;1481del:93
- 1473del:93;1474del:93;1475del:93;1476del:93;1477del:93;1478del:93;1479del:93;1480del:93;1481del:93
- 1473del:93;1474del:93;1475del:93;1476del:93;1477del:93;1478del:93;1479del:93;1480del:93;1481del:93

(ORF ends at 1470)

PacRAT: `c.1466_1470del = p.Pro489fs`

- 1258del;1466del;1467del;1468del;1469del;1470del
- 737del;937del;1011del;1160del;1439_1440insT;1466del;1467del;1468del;1469del;1470del
- 1466del;1467del;1468del;1469del;1470del
- 1466del;1467del;1468del;1469del;1470del
- 1466del;1467del;1468del;1469del;1470del
- 19del;122del;123del;177_178insG;521_522insG;1258del;1466del;1467del;1468del;1469del;1470del
- 10del;19del;168del;295_296insT;775del;876_877insA;1061_1062insC;1126del;1286_1287insA;1366del;1431_1432insG;1435_1436insT;1466del;1467del;1468del;1469del;1470del

Again, disagreement on where the deletion happens. Pacybara places it outside of the ORF

#### Barcode GGATTCATAAACGAACTA

Pacybara: `c.[665T>G;1446_1449delinsG] = p.[Ile222Ser;Phe482_Tyr483del]`

- -2del:93;665T>G:93;1446del:93;1447del:93;1448del:93;1449del:93;1448A>G:93
- 665T>G:93;1446del:87;1447del:87;1448del:87;1449del:87;1448A>G:93;1592del:93;1598G>A:68
- 665T>G:93;1258del:50;1446del:93;1447del:93;1448del:93;1449del:93;1448A>G:93
- 665T>G:93;927A>C:17;1446del:93;1447del:93;1448del:93;1449del:93;1448A>G:92
- 665T>G:93;1446del:93;1447del:93;1448del:93;1449del:93;1448A>G:93
- 338del:66;665T>G:93;1446del:93;1447del:93;1448del:93;1449del:93;1448A>G:93
- -2del:50;665T>G:93;1258del:63;1393del:53;1446del:93;1447del:93;1448del:93;1449del:93;1448A>G:93;1592del:49
- 665T>G:93;1293insA:93;1446del:93;1447del:93;1448del:93;1449del:93;1448A>G:93
- 665T>G:93;1258del:52;1446del:93;1447del:93;1448del:93;1449del:93;1448A>G:93

PacRAT: `c.[665T>G;1443_1448delinsTG] = p.Pro481fs`

- 665T>G;1258del;1443del;1444del;1445del;1446del;1448A>G
- 665T>G;1443del;1444del;1445del;1446del;1448A>G
- 338del;665T>G;1443del;1444del;1445del;1446del;1448A>G
- 665T>G;1258del;1393del;1443del;1444del;1445del;1446del;1448A>G
- 665T>G;1258del;1443del;1444del;1445del;1446del;1448A>G
- 665T>G;1443del;1444del;1445del;1446del;1448A>G
- 665T>G;1443del;1444del;1445del;1446del;1448A>G
- 665T>G;927A>C;1443del;1444del;1445del;1446del;1448A>G
- 665T>G;1292_1293insA;1443del;1444del;1445del;1446del;1448A>G
- 19del;27G>T;220del;253_254insA;530del;665T>G;669_670insG;686del;691A>C;721del;763del;767_768insC;844del;860del;866A>T;867T>A;868A>T;921del;945_946insC;994del;1006T>C;1007C>T;1125_1126insT;1139del;1219del;1258_1259insC;1303T>G;1443del;1444del;1445del;1446del;1448A>G

Again, the positioning of the indel is off.

### Supplemental Text 2: Non-unique barcodes in *LDLR* : Pacybara vs PacRAT

Here we manually examined the read information underlying variant calls for 8 arbitrarily picked barcodes from the *LDLR* library for which PacRAT and Pacybara disagreed and which were considered non-unique. We found 6 out of 8 of these cases to be correctly handled by Pacybara. One of them was better handled by PacRAT, and the one remaining case was undecidable.

#### Barcode CAGACTGTCTCAGTGAGAGAGTCTC

Pacybara call: Non-unique barcode, with two clusters: `c.[509_510insC;2591del] = p.Asp170fs` and `WT`, respectively

PacRAT call: `WT`

Divergent basecalls in reads used by Pacybara:

Cluster 1:

- `-154del(Q93);509_510insC(Q93);2591del(Q93)`
- `2591del(Q93)`
- `509_510insC(Q6);2591del(Q93)`

Cluster 2:

- `WT`

The same four reads were used by PacRAT.

The reads speak more in favor of Pacybara here.

#### Barcode GAGTCACTGAGACTCTCAGTGTGTC

Pacybara call: Non-unique barcode, with two clusters: `c.278T>A = p.Val93Glu` and `c.262_264delinsTTT = p.Arg88Phe`

PacRAT call: `WT`

Divergent basecalls in reads used by Pacybara:

Cluster 1:

- `278T>A(Q93)`
- `278T>A(Q93)`

Cluster 2:

- `262A>T(Q93);263G>T(Q93);264G>T(Q93)`

PacRAT admitted 8 reads for this barcode, (5 of which were filtered out by Pacybara)

- `278T>A`
- `278T>A`
- `262A>T;263G>T;264G>T`
- `=`
- `319del;320A>C`
- `56_57insG;93_94insT;278T>A;465_466insA;645_646insC;645_646insG;654_655insT;654_655insG;656G>C;703_704insC;763_764insC;934_935insA;1228_1229insG;1268_1269insC;1311_1312insA;1360_1361insC;1539_1540insA;1606_1607insG;1819_1820insC;1819_1820insA;1820A>C;1938_1939insC;1938_1939insC;1938_1939insT;1942_1943insC;2069A>C;2070C>T;2071T>C;2171_2172insA;2304_2305insC`
- `190del;278T>A;946del;1607G>T;1704del;1705del;1706del;1764_1765insA;1766_1767insC;1939_1940insT;1939_1940insA;2051del;2064del;2114del;2218del;2219del;2249del`
- `176del`

Of the reads PacRAT considered, `c.278T>A` occurred in 4 out of 8, but was discarded from the consensus.

The read evidence speaks more in favor of Pacybara here.

#### Barcode CACTCACTGACACACTGAGTGAGAC

Pacybara call: Non-unique barcode, with two clusters: `c.507C>T        = p.Asn169=` and `c.507_508insT = p.Asn169fs`

PacRAT call: `c.507_508insT = p.Asn169fs`

Divergent base-calls in reads used by Pacybara:

Cluster 1:

- -178del(Q93);507C>T(Q93)
- 507C>T(Q93);2063_2064insC(Q54)
- 507C>T(Q93)
- -177_-178insA(Q7);507C>T(Q93);1360_1361insC(Q5)
- 209del(Q93);507C>T(Q93);1157_1158insC(Q93)
- 507C>T(Q93);2619+163G>A(Q93)
- 507C>T(Q93)

Cluster 2:

- 450_451insA(Q93);507_508insT(Q78)
- 507_508insT(Q58)

PacRAT admitted 17 reads for this barcode (8 of which were filtered out by Pacybara)

- 504_505insA;1424del
- 292G>C;293G>A;294C>T;509_510insC;1942_1943insC
- 507C>T
- 507C>T
- 507C>T;1360_1361insC
- 507C>T
- 507C>T
- 450_451insA;507_508insT
- 1405A>G;1817del;1906del
- 507_508insT
- 209del;507C>T;1157_1158insC
- 507_508insT
- 507_508insT
- 507C>T;509_510insC;974_975insT;1019_1020insC;1081del;1304_1305insG;1633del;1849del;2079_2080insT;2278_2279insC;2340T>G;2350del;2539_2540insC
- 507C>T

- 22del;56_57insG;56_57insG;61A>G;62C>G;94T>G;96C>T;127A>T;129G>T;209C>T;210C>G;238A>G;239A>C;240C>G;276A>T;309_310insT;324G>T;370C>A;371G>C;406G>T;408C>T;433G>T;434T>G;455del;2116_2117insG
- 507C>T;2063_2064insC

The results appear more in favor if Pacybara, even though there aren't technically two clusters here. Just a frequent sequencing error that misrepresents a substitution as an indel.

#### Barcode CAGTGTGTCTGTGTGTGAGAGACAC

Pacybara call: Non-unique barcode, with two clusters: `c.1997G>A = p.Trp666Ter` and `WT`

PacRAT call: `WT`

Divergent basecalls in reads used by Pacybara:

Cluster 1:

- 1997G>A(Q93)
- 1997G>A(Q93)
- 1997G>A(Q93)
- 1997G>A(Q93)

Cluster 2:

- WT
- 1744_1745insG(Q34);2397_2398insG(Q93);2619+6del(Q93)

PacRAT extracted 11 reads for this barcode (5 of which were filtered out by Pacybara)

- 1997G>A
- 1997G>A
- WT
- 269_270insT;274C>A;275A>C;276A>C;505_506insA;505_506insC;505_506insG;1207_1208insC;1398_1399insA;2227_2228insC
- 1997G>A
- 1485del;1997G>A;2105_2106insG
- WT
- 1997G>A
- 509_510insC
- 1744_1745insG;2397_2398insG
- 553_554insG;1997G>A

The evidence appears to be in favor of Pacybara again.

#### Barcode GTGAGAGAGTCAGACAGAGTGAGAG

Pacybara call: Non-unique barcode, with two clusters: `c.1480G>A = p.Val494Ile` and `WT`

PacRAT call: WT

Divergent basecalls in reads used by Pacybara:

Cluster 1:

- 1480G>A(Q93)
- 1480G>A(Q32)
- 1480G>A(Q93)

Cluster 2:

- WT

PacRAT extracted 10 reads for this barcode (6 of which were filtered out by Pacybara)

- WT
- 1480G>A
- 1480G>A
- 416A>G;417C>A
- 31_32insC;509_510insC;2093_2094insC
- 57del;553_554insG;574del;737_738insA;2046del
- WT
- WT
- 1480G>A
- 222_223insT;225_226insG;312_313insC;447_448insC;504_505insA;656_657insC;666_667insA;672_673insA;833_834insA;865_866insG;917_918insA;917_918insG;924_925insC;960_961insA;976_977insC;1032_1033insT;1157_1158insC;1221_1222insG;1323_1324insT;1396_1397insA;1396_1397insC;1411_1412insG;1421_1422insG;1492_1493insT;1492_1493insC;1524_1525insA;1632_1633insA;1632_1633insG;1656_1657insT;1773_1774insG;1814_1815insG;1820_1821insC;1839_1840insT;1992_1993insA;2122_2123insT;2133_2134insC;2133_2134insT;2225_2226insC;2225_2226insA;2238_2239insC;2244_2245insA;2293_2294insT;2328_2329insA;2328_2329insG;2336_2337insA;2346_2347insA;2381_2382insC;2433_2434insA;2483_2484insT;2507_2508insC;2507_2508insG;2516_2517insC;2540_2541insC

The evidence appears to be in favor of Pacybara again.

#### Barcode CTGACTGTCTGTGAGACAGACAGAG

Pacybara call: Non-unique barcode, with two clusters: c.[446G>A;1157_1158insC] = p.[Gly149Asp;Asp386fs] and WT

PacRAT call: c.446G>A = p.Gly149Asp

Divergent basecalls in reads used by Pacybara:

Cluster 1:

- 446G>A(Q93);1157_1158insC(Q7)
- 446G>A(Q93)
- 446G>A(Q93);1157_1158insC(Q93)

Cluster 2:
- 645_646insT(Q6);2619+7_8insC(Q93)

PacRAT picked admitted 10 reads (6 of which were filtered by Pacybara)

- 446G>A
- 446_447insA;1268T>A
- 446_447insA
- 394_395insG;446G>A;452del;1511_1512insG;1876_1877insA;1930_1931insA
- 446G>A
- 384_385insG;446G>A;654_655insG;781_782insG;844T>G;2025_2026insG;2025_2026insG;2025_2026insC;2025_2026insC;2025_2026insA;2025_2026insA;2025_2026insT;2402_2403insC
- 446G>A;1157_1158insC
- 446G>A
- 645_646insT
- 446G>A;2550A>G

Here, Pacybara gets it wrong by including a variant that's likely spurious. c.1157_1158insC also appears to be a frequent flyer error seen in many unrelated clusters.

### Barcode CTGACAGACACTCAGACTCTCTCTG

Pacybara call: Non-unique barcode, with two clusters: `c.[364_366delinsCTG;457_459delinsAAG] = p.[Ile122Leu;Phe153Lys]` and `c.[447_448insC;1380C>G] = p.Gly149fs`

PacRAT call: `WT`

Divergent basecalls in reads used by Pacybara:

Cluster 1:

- `-177_-178insA(Q93);364A>C(Q79);366C>G(Q93);457T>A(Q93);458T>A(Q50);459C>G(Q93)`
- `364A>C(Q93);366C>G(Q93);457T>A(Q16);458T>A(Q31);459C>G(Q57);2592del(Q93)`

Cluster 2:

- `-150_-151insC(Q93);447_448insC(Q93);1380C>G(Q93)`

PacRAT admitted 4 reads for this barcode (1 of which was filtered out by Pacybara)

- 364A>C;366C>G;457T>A;458T>A;459C>G
- 364A>C;366C>G;457T>A;458T>A;459C>G
- 447_448insC;1380C>G
- 10_11insG;31_32insC;68_69insG;136_137insG;225_226insG;225_226insG;234_235insG;312_313insC;328_329insC;482_483insC;520G>A;522A>T;656_657insC;735_736insG;823_824insT;939_940insG;976_977insC;1016_1017insG;1025_1026insC;1184_1185insG;1399_1400insC;1423_1424insC;1721_1722insC;1810_1811insG;2083_2084insC;2266_2267insC;2302_2303insC;2408_2409insC;2510_2511insC

The evidence appears to be in favor of Pacybara again.

#### Barcode CTCAGTGAGTGACTGAGTCTGTCAG

Pacybara call: Non-unique barcode, with two clusters: `WT` and `c.553A>C = p.Arg185=`

PacRAT call: `c.553_554delinsCT = p.Arg185Leu`

Divergent basecalls in reads used by Pacybara:

Cluster 1:

- -51_-52insG(Q93);552_553insC(Q93);553A>T(Q12)
- -51_-52insG(Q22);220A>C(Q93);221G>T(Q93);222C>T(Q93);289A>C(Q93);290A>C(Q93);291C>G(Q76);1485del(Q48)

Cluster 2:

- 527G>T(Q89);553A>C(Q93);554G>T(Q76)

PacRAT extracted 7 reads; might have a correct inkling here.

- 552_553insC;553A>T
- 554G>T
- 553A>C
- 553A>C;554G>T;922_923insA
- 220A>C;221G>T;222C>T;289A>C;290A>C;291C>G;1485del
- 40T>A;56_57insG;87del;168T>A;224_225insC;224_225insT;225T>G;340del;503_504insG;503_504insC;553del;554G>T;584_585insC;651del;830A>T;1108del;1268_1269insC;1282del;1306G>T;1307T>G;1320del;1606_1607insG;1649_1650insG;2212del;2221_2222insC
- 527G>T;553A>C;554G>T

Hard to say whether either tool got it right here. Lots of confusing indel errors are complicating this call.

Interestingly, Pacybara's clustering decision was made here due to an unrelated variant outside of the ORF (`c.-51_-52insG`)

### Supplemental Tables

| uptags associated with: | single downtag | multiple downtags |
|---|---|---|
| single genotype | 20,125 | 58,703 |
| multiple genotype | 1 | 15,658 |

Table S1: Non-unique barcodes in the LDLR library when enabling or disabling virtual barcodes. Virtual barcodes also detect differences in downtags, which help resolve PCR chimeras

| **uptag seq** | **downtag seq** | **reads** | **ORF genotype** | **off-target mut.** |
|---|---|---|---|---|
| AACACTGAGTCAGAGACACACAGTG | CTCACAGTGTGACTCTCACACAGAG | 1 | c.544C>G | NA |
| AACACTGAGTCAGAGACACACAGTG | CTCTCACTGACAGAGTGTCACAGAC | 1 | c.544C>G | NA |
| ACACAGTCTCTGTGACACTGTGTC | CTGTCACTCAGACTCAGTCACTCTC | 8 | c.290_291delinsTG | -32C>G |
| ACACAGTCTCTGTGACACTGTGTC | GACTGAGAGAGTCTGTGTCTGACTG | 1 | c.290_291delinsTG | -32C>G |
| ACACAGTCTCTGTGACACTGTGTC | CTCACACACAGCTCTGCTCGACTGTGAC | 1 | c.[290_291delinsTG;554_555delinsTT] | -32C>G |
| ACACAGTCTCTGTGACACTGTGTC | CTCTGACACTCACTGACAGTCTCAG | 1 | c.[290_291delinsTG;2580G>A] | -32C>G |
| ACACAGTCTCTGTGACACTGTGTC | GCTGTCACTCAGACTCAGTCTCTG | 1 | c.290_291delinsTG | -32C>G |
| ACACAGTCTCTGTGACACTGTGTC | CTGTCACTCAGACTCAGTCACTCTC | 1 | c.= | NA |
| ACACAGTCTCTGTGACACTGTGTC | CAGTCACTCACGACTCAGTCACTCTC | 1 | c.290_291delinsTG | -32C>G |
| ACTCACTGTGAGTGTGAGTGTGAG | CTCTGAGTGAGACAGTCTGTGTCTG | 1 | c.234T>G | NA |
| ACTCACTGTGAGTGTGAGTGTGAG | CACTCTCAGACTGTCACTCTGACTC | 1 | c.234T>G | NA |
| ACTCACTGTGAGTGTGAGTGTGAG | CTCTGAGTGAGACAGTCTGTGTCTG | 1 | c.234T>G | NA |

| | | | | |
|---|---|---|---|---|
| ACTCACTGTGAGTGTGAGTGTGAG | CTCTGAGTGAGACAGTCTGTGTCTG | 1 | c.234T>G | NA |
| ACTCACTGTGAGTGTGAGTGTGAG | CACTGACTGTGACTGTCAGTCTGTC | 1 | c.364_366delinsCCG | NA |
| ACTCACTGTGAGTGTGAGTGTGAG | CTCTGAGTGAGACAGTCTGTGTCTG | 1 | c.234T>G | NA |
| ACTCACTGTGAGTGTGAGTGTGAG | CTCACTGACACTCTGTGTGTCTGAG | 1 | c.234T>G | NA |
| ACTCACTGTGAGTGTGAGTGTGAG | CTCTGAGTGAGACAGTCTGTGTCTG | 1 | c.[234T>G;1559del] | NA |
| ACTCACTGTGAGTGTGAGTGTGAG | CTCTGAGTGAGACAGTCTGTGTCTG | 1 | c.234T>G | NA |
| ACTCACTGTGAGTGTGAGTGTGAG | CTCTGAGTGAGACAGTCTGTGTCTG | 1 | c.234T>G | NA |
| ACTCACTGTGAGTGTGAGTGTGAG | CTCTGAGTGAGACAGTCTGTGTCTG | 1 | c.234T>G | NA |
| ACTCACTGTGAGTGTGAGTGTGAG | CTCTGAGTGAGACAGTCTGTGTCTG | 1 | c.234T>G | NA |
| ACAGACACTCTGTGTCTCTGTGTC | CCCAGACACACAGAGACTCTCTGTC | 4 | c.[345C>T;377_378delinsGG] | NA |
| ACAGACACTCTGTGTCTCTGTGTC | CACAGAGAGTGTGACACTGACTGTG | 1 | c.[345C>T;377_378delinsGG] | NA |
| ACAGACACTCTGTGTCTCTGTGTC | GACTCACAGAGAGAGACACACACAG | 1 | c.[345C>T;377_378delinsGG] | NA |
| ACAGACACTCTGTGTCTCTGTGTC | CACAGTCACTGAGCTGACGCAGTCTC | 1 | c.[345C>T;377_378delinsGG] | NA |

Table S2: Examples of chimeras detected in the *LDLR* library using upstream and downstream barcodes.
